# Linking the International Wheat Genome Sequencing Consortium bread wheat reference genome sequence to wheat genetic and phenomic data

**DOI:** 10.1101/363259

**Authors:** Michael Alaux, Jane Rogers, Thomas Letellier, Raphaël Flores, Françoise Alfama, Cyril Pommier, Nacer Mohellibi, Sophie Durand, Erik Kimmel, Célia Michotey, Claire Guerche, Mikaél Loaec, Mathilde Lainé, Delphine Steinbach, Frédéric Choulet, Hélène Rimbert, Philippe Leroy, Nicolas Guilhot, Jérôme Salse, Catherine Feuillet, International Wheat Genome Sequencing Consortium, Etienne Paux, Kellye Eversole, Anne-Françoise Adam-Blondon, Hadi Quesneville

## Abstract

The Wheat@URGI portal (https://wheat-urgi.versailles.inra.fr) has been developed to provide the international community of researchers and breeders with access to the bread wheat reference genome sequence produced by the International Wheat Genome Sequencing Consortium. Genome browsers, BLAST, and InterMine tools have been established for in depth exploration of the genome sequence together with additional linked datasets including physical maps, sequence variations, gene expression, and genetic and phenomic data from other international collaborative projects already stored in the GnpIS information system. The portal provides enhanced search and browser features that will facilitate the deployment of the latest genomics resources in wheat improvement.

## Background

The International Wheat Genome Sequencing Consortium (IWGSC) [1] is an international collaborative group of growers, academic scientists, and public and private breeders that was established to generate a high quality reference genome sequence of the hexaploid bread wheat, and to provide breeders with state-of-the-art tools for wheat improvement. The vision of the consortium is that the high quality, annotated ordered genome sequence integrated with physical maps will serve as a foundation for the accelerated development of improved varieties and will empower all aspects of basic and applied wheat science to address the important challenge of food security. A first analysis of the reference sequence produced by the consortium (IWGSC RefSeq v1.0) was recently published [2].

To ensure that wheat breeding and research programs can make the most of this extensive genomic resource, the IWGSC endorsed the establishment of a data repository at URGI (Unité de Recherche Génomique Info / research unit in genomics and bioinformatics) from INRA (Institut National de la Recherche Agronomique / French national institute for agricultural research) to develop databases and browsers with relevant links to public data available worldwide. The IWGSC data repository is thus hosted by URGI to support public and private parties in data management as well as analysis and usage of the sequence data. Wheat functional genomics (expression, methylation, etc.), genetic, and phenomic data has increased concurrently, requiring the development of additional tools and resources to integrate different data for biologists and breeders. To manage this escalation of data, URGI have built this data repository for the wheat community with the following specific aims: (i) store resources for which no public archive exists (e.g. physical maps, phenotype information); (ii) enable pre-publication access to specific datasets (e.g. sequence assemblies and annotations, physical maps, markers); and (iii) rapid release of integrated resources upon publication. The repository has been designed in accordance with the “FAIR” principles [3] to ensure that the data are Findable, Accessible, Interoperable and Reusable. To address the challenge of integrating diverse data types from multiple sources, URGI employs solutions that provide enhanced features for data exploration, mining and visualisation using the GnpIS information system [4] combined with a high level of data interoperability.

Here we describe the data and tools currently available through the Wheat@URGI portal [5], the primary resource for the reference sequence of the bread wheat genome (IWGSC RefSeq v1.0) and other IWGSC wheat genomic data. The links to functional genomics, genetic and phenomic data from many other large wheat projects are also described.

## A large wealth of data is available throughout the Wheat@URGI portal

The data hosted by the Wheat@URGI portal are available through flat files stored in the IWGSC data repository and through the GnpIS information system [4]. GnpIS encompasses a set of integrated databases to manage genomic data using well-known tools such as BLAST, JBrowse, GBrowse and InterMine, and an in-house database called GnpIS-coreDB developed by URGI to manage genetic and phenomic data.

### IWGSC data

Through its concerted efforts to achieve a high quality, functionally annotated reference wheat genome sequence, the IWGSC has developed a variety of resources for the bread wheat *(Triticum aestivum L.)* accession Chinese Spring. The IWGSC data hosted in the Wheat@URGI portal within the IWGSC data repository are shown in Table 1. They fall into four broad categories: (i) physical maps, (ii) sequence assemblies and annotations, (iii) gene expression, and (iv) variation data.

**Table 1.**
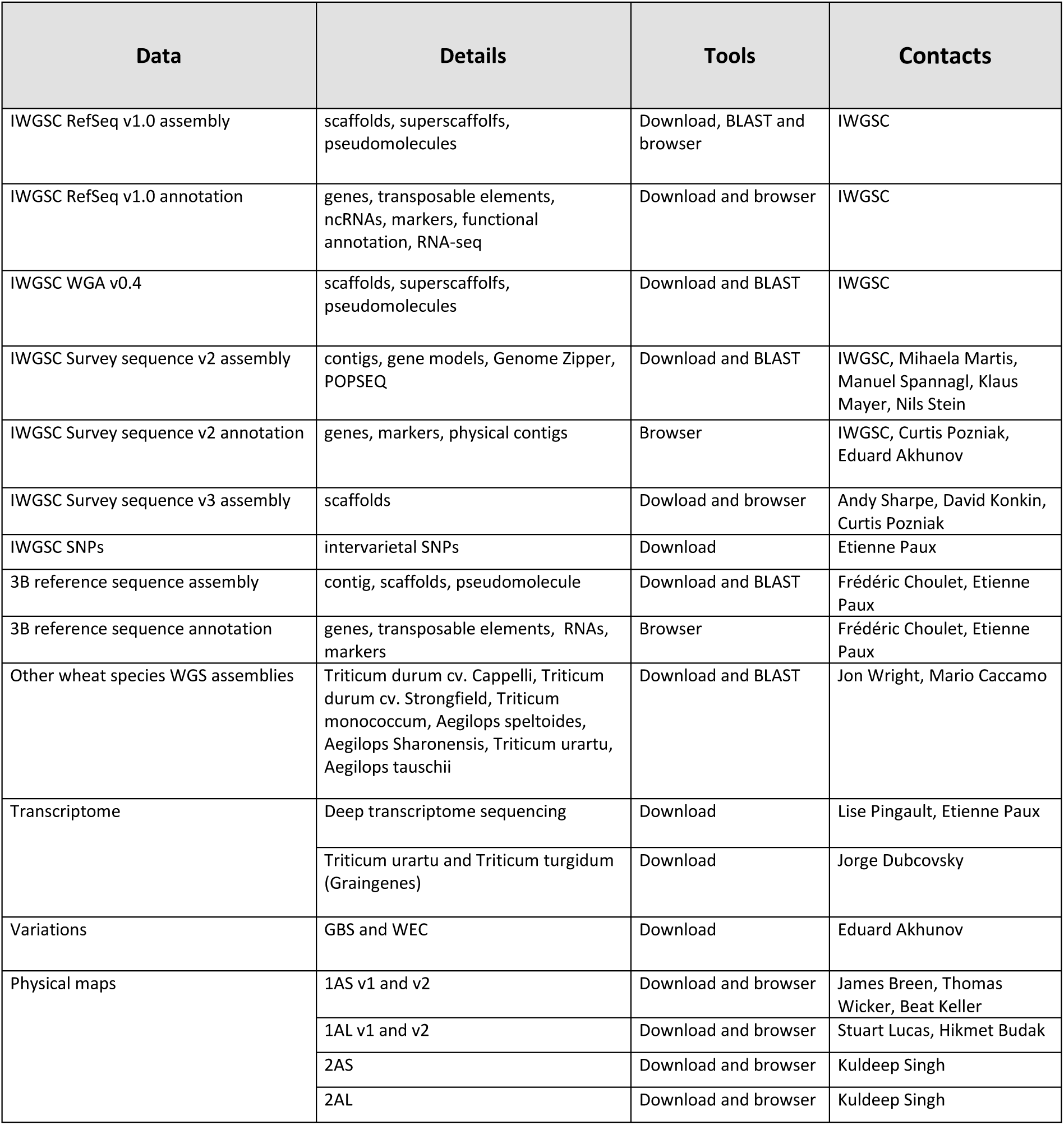
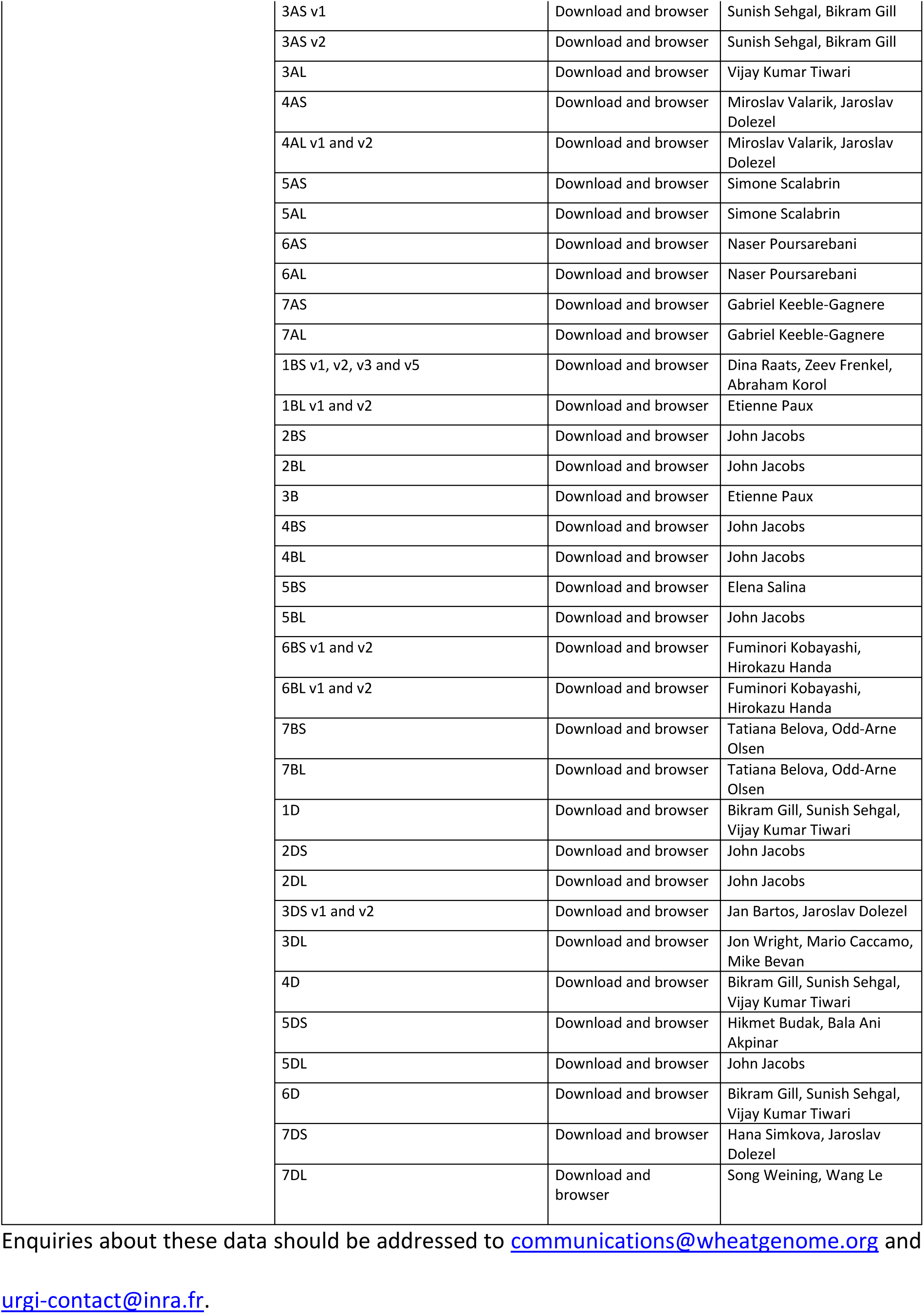
IWGSC data summary in open access hosted in the IWGSC Data Repository of the Wheat@URGI portal in March 2018.

| Data | Details | Tools | Contacts |
| --- | --- | --- | --- |
| IWGSC RefSeq v1.0 assembly | scaffolds, superscaffolds, pseudomolecules | Download, BLAST and browser | IWGSC |
| IWGSC RefSeq v1.0 annotation | genes, transposable elements, ncRNAs, markers, functional annotation, RNA-seq | Download and browser | IWGSC |
| IWGSC WGA v0.4 | scaffolds, superscaffolds, pseudomolecules | Download and BLAST | IWGSC |
| IWGSC Survey sequence v2 assembly | contigs, gene models, Genome Zipper, POPSEQ | Download and BLAST | IWGSC, Mihaela Martis, Manuel Spannagl, Klaus Mayer, Nils Stein |
| IWGSC Survey sequence v2 annotation | genes, markers, physical contigs | Browser | IWGSC, Curtis Pozniak, Eduard Akhunov |
| IWGSC Survey sequence v3 assembly | scaffolds | Download and browser | Andy Sharpe, David Konkin, Curtis Pozniak |
| IWGSC SNPs | intervarietal SNPs | Download | Etienne Paux |
| 3B reference sequence assembly | contig, scaffolds, pseudomolecule | Download and BLAST | Frédéric Choulet, Etienne Paux |
| 3B reference sequence annotation | genes, transposable elements, RNAs, markers | Browser | Frédéric Choulet, Etienne Paux |
| Other wheat species WGS assemblies | Triticum durum cv. Cappelli, Triticum durum cv. Strongfield, Triticum monococcum, Aegilops speltoides, Aegilops Sharonensis, Triticum urartu, Aegilops tauschii | Download and BLAST | Jon Wright, Mario Caccamo |
| Transcriptome | Deep transcriptome sequencing | Download | Lise Pingault, Etienne Paux |
|  | Triticum urartu and Triticum turgidum (Graingenes) | Download | Jorge Dubcovsky |
| Variations | GBS and WEC | Download | Eduard Akhunov |
| Physical maps | 1AS v1 and v2 | Download and browser | James Breen, Thomas Wicker, Beat Keller |
|  | 1AL v1 and v2 | Download and browser | Stuart Lucas, Hikmet Budak |
|  | 2AS | Download and browser | Kuldeep Singh |
|  | 2AL | Download and browser | Kuldeep Singh |

|  |  |  |
| --- | --- | --- |
| 3AS v1 | Download and browser | Sunish Sehgal, Bikram Gill |
| 3AS v2 | Download and browser | Sunish Sehgal, Bikram Gill |
| 3AL | Download and browser | Vijay Kumar Tiwari |
| 4AS | Download and browser | Miroslav Valarik, Jaroslav Dolezel |
| 4AL v1 and v2 | Download and browser | Miroslav Valarik, Jaroslav Dolezel |
| 5AS | Download and browser | Simone Scalabrin |
| 5AL | Download and browser | Simone Scalabrin |
| 6AS | Download and browser | Naser Poursarebani |
| 6AL | Download and browser | Naser Poursarebani |
| 7AS | Download and browser | Gabriel Keeble-Gagnere |
| 7AL | Download and browser | Gabriel Keeble-Gagnere |
| 1BS v1, v2, v3 and v5 | Download and browser | Dina Raats, Zeev Frenkel, Abraham Korol |
| 1BL v1 and v2 | Download and browser | Etienne Paux |
| 2BS | Download and browser | John Jacobs |
| 2BL | Download and browser | John Jacobs |
| 3B | Download and browser | Etienne Paux |
| 4BS | Download and browser | John Jacobs |
| 4BL | Download and browser | John Jacobs |
| 5BS | Download and browser | Elena Salina |
| 5BL | Download and browser | John Jacobs |
| 6BS v1 and v2 | Download and browser | Fuminori Kobayashi, Hirokazu Handa |
| 6BL v1 and v2 | Download and browser | Fuminori Kobayashi, Hirokazu Handa |
| 7BS | Download and browser | Tatiana Belova, Odd-Arne Olsen |
| 7BL | Download and browser | Tatiana Belova, Odd-Arne Olsen |
| 1D | Download and browser | Bikram Gill, Sunish Sehgal, Vijay Kumar Tiwari |
| 2DS | Download and browser | John Jacobs |
| 2DL | Download and browser | John Jacobs |
| 3DS v1 and v2 | Download and browser | Jan Bartos, Jaroslav Dolezel |
| 3DL | Download and browser | Jon Wright, Mario Caccamo, Mike Bevan |
| 4D | Download and browser | Bikram Gill, Sunish Sehgal, Vijay Kumar Tiwari |
| 5DS | Download and browser | Hikmet Budak, Bala Ani Akpinar |
| 5DL | Download and browser | John Jacobs |
| 6D | Download and browser | Bikram Gill, Sunish Sehgal, Vijay Kumar Tiwari |
| 7DS | Download and browser | Hana Simkova, Jaroslav Dolezel |
| 7DL | Download and browser | Song Weining, Wang Le |

#### Physical maps

physical maps assembled by IWGSC scientists for the 21 bread wheat chromosomes, based on high information content fluorescence fingerprinting (HICF) [6] or whole genome profiling (WGP^™^) [7] of flow-sorted chromosome or chromosome-arm specific BAC libraries, are stored and displayed. The positions of individual BAC clones, markers, and deletion bins are mapped onto physical contigs. The database maintains all released versions of each physical map with the software used to produce the BAC clone assemblies (FPC [8] or LTC [9]), information from the group that produced the map and a link to order the BAC clones from the French plant genomic resource centre [10].

#### Sequence assemblies and annotations

the IWGSC wheat genome sequence assemblies available for download, BLAST [11], and display in genome browsers include the draft survey sequence assemblies released in 2014 (IWGSC Chromosome Survey Sequence (CSS) v1) and two improved versions (CSS v2 and v3) [12], and the chromosome 3B reference sequence (the first reference quality chromosome sequence obtained by the consortium) [13]. Associated with these assemblies are the virtual gene order map generated for the CSS (Genome Zipper), the POPSEQ data used to order sequence contigs on chromosomes [14] and mapped marker sets. The reference sequence of the bread wheat genome (IWGSC RefSeq v1.0, 14.5 Gb assembly with super scaffold N50 of 22.8Mb) was obtained by integrating whole genome shotgun Illumina short reads assembled with NRGene’s DeNovoMAGIC^™^ software with the wealth of IWGSC map and sequence resources [2]. The IWGSC RefSeq v1.0 is available for download, BLAST, and browser display. Users can access the whole genome, pseudomolecules of individual chromosomes or chromosome arms, and scaffolds with the structural and functional annotation of genes, transposable elements, and non-coding RNAs generated by the IWGSC. In addition, mapped markers as well as alignments of nucleic acid and protein evidence supporting the annotation are available. Updated versions of the annotation for genes belonging to specific gene families or regions of specific chromosomes that have been manually annotated (ca. 3685 genes) can be found in the IWGSC RefSeq v1.1 annotation.

In addition to the bread wheat sequence, the IWGSC also assembled seven diploid and tetraploid wheat related species: *Triticum durum* cv. Cappelli, *Triticum durum* cv. Strongfield, *Triticum durum* cv Svevo, *Triticum monococcum*, *Triticum urartu*, *Aegilops speltoides*, *Aegilops sharonensis* [12]. Download and BLAST is available for these data.

#### Expression data

RNA-Seq expression data are available as reads counts and transcripts per kilobase million (TPM) for the IWGSC RefSeq v1.1 annotation. It is a transcriptome atlas developed from 850 RNA-Seq datasets representing a diverse range of tissues, development stages and environmental condition [15].

#### Variation data

These data consists of downloadable VCF files from genotyping by sequencing and whole exome capture experiments of 62 diverse wheat lines [16] and of the IWGSC 3,289,847 inter-varietal SNPs [17].

### Wheat gene pool

As well as IWGSC resources, URGI also hosts other open access wheat sequence data to facilitate research into the wheat gene pool. Sequence assemblies available for download and BLAST include the bread wheat whole genome sequence assembly *Triticum aestivum* TGACv1 [18] and the diploid progenitor of *Aegilops tauschii* [19].

### Genetic and phenomic data

In addition to sequence data, the Wheat@URGI portal hosts, within GnpIS-coreDB, several sets of genetic and phenomic wheat data [20] that have been produced from French, European, and international projects since 2000 [21]. A significant amount of these data is available without restriction. However, access to restricted data can be obtained through a material transfer or intellectual property agreement. Table 2 presents the types and number of genetic and phenomic data hosted in the GnpIS-coreDB database.

**Table 2.** Genetic and phenomic wheat data summary hosted in the GnpIS-coreDB database of the Wheat@URGI portal in March 2018.

| Data type | Object | #Total | #Open access | #Restricted access to projects |
| --- | --- | --- | --- | --- |
| Genetic Resources | Taxon | 56 | 56 | 0 |
|  | Accession | 12839 | 10016 | 2823 |
| Genetic Maps | Map | 30 | 29 | 1 |
|  | Marker | 704822 | 34164 | 670658 |
|  | QTL | 749 | 465 | 284 |
| SNP discovery | Sequence Variation | 4189312581 | 90 | 4189312491 |
|  | SNP, indel | 724132 | 95 | 724037 |
| Genotyping (high throughput) | Experiment | 23 | 2 | 21 |
|  | Sample | 8885 | 47 | 8872 |
|  | Marker | 668540 | 0 | 668540 |
| Phenotyping | Trial | 850 | 821 | 29 |
|  | Plot | 3660 | 2985 | 901 |
|  | Variable | 282 | 89 | 195 |
|  | Observation | 1171172 | 527981 | 643191 |
| GWAS | Analysis | 1555 | 43 | 1512 |
|  | Sample | 2365 | 1839 | 526 |
|  | Variable | 359 | 37 | 322 |
|  | Marker | 123866 | 4109 | 119757 |
|  | Association | 824217 | 48596 | 775621 |

Genetic information corresponds to genetically mapped markers, quantitative trait loci (QTLs), genetic resources (germplasms), and genetic studies (genome wide association studies - GWAS). Genomic information consists of variation from SNP discovery experiments, genotyping, comparative genomics (synteny) and expression data (microarray, RNA-Seq). Phenomic data are available as whole trials including phenotypic and environmental observations recorded using ontologies controlled variables with MIAPPE [22] compliant metadata.

Germplasm data were mainly provided by the French small grain cereals genebank maintained by INRA at Clermont-Ferrand [23] but also by partners of several EU projects. They were linked together with related genotyping or phenotyping characterization data. Generally, genetic and phenomic data have been produced by INRA and its partners in large collaborative projects.

## Browsing and searching a large variety of integrated data

Data can be easily accessed through the Wheat@URGI portal [5] using (i) tabs at the top of the pages allowing access in one click to the data, tools, and projects descriptions as well as the IWGSC data repository, (ii) direct links on the home page to the different data types (e.g. clicking on “Physical maps” opens the physical maps browser), and (iii) data discovery and InterMine [24] tools on the home page.

The IWGSC data repository [25] allows accessing consortium data by (i) clicking on a chromosome to open a pop-up menu with all related data (*e.g.* 3A, 3B, etc.), (ii) using the tabs on the left to access the data by type (e.g. Assemblies, Annotations, etc.), or useful links to the news, the BLAST tool, the FAQ, the access status of the data (e.g. open access), etc.

### Physical maps browser

A GBrowse [26] displays the physical maps generated by the IWGSC members [27]. A clickable image on the top of the browser gives access to all versions of the physical map for each chromosome. The browser displays physical contigs, BACs, deletions bins, and markers. From the BACs track, it is possible to order BAC clones directly at the INRA French plant genomic resource centre [10]. From the BACs and markers tracks, one can go directly to the corresponding region in the IWGSC RefSeq v1.0 browser.

### Genome browser and BLAST

The IWGSC RefSeq v1.0 is displayed in a dedicated JBrowse [28], [29]. The “markers track” provides links to additional genetic information stored in GnpIS-coreDB which includes access to the position of the marker in cM on genetic maps and to the overlapping QTLs. The most popular tool of the IWGSC data repository is the BLAST search tool (476,000 BLAST searches launched in 2017). All of the wheat sequences available on the Wheat@URGI portal are indexed for BLAST search (see [30] for the complete list). A set of databanks can be selected: e.g. IWGSC RefSeq v1.0 and IWGSC CSS v3 for a given chromosome. The result is presented in a classical tabular format with (i) links to download the data (matching contigs and high scoring pairs - HSP), (ii) links on the genome browsers directly zooming in on the matching region and (iii) external links to EnsemblPlants [31].

### Genetic and phenomic data in GnpIS-coreDB

The IWGSC sequence data are linked to genetic and phenomic data within the GnpIS information system [4]. This integration is organized around key data, also called “pivot data” as they are pivotal objects which allow integration between data types. The key objects used to link genomic resources to genetic data are markers and QTLs. Markers are mapped on the genome sequences and provides information on neighbour genes and their function. They also have links to GnpIS-coreDB genetic maps, QTLs, genotyping and GWAS data. Additional information on the marker itself can be found regarding the marker type (e.g. SSR, DArT), the primers sequence for PCR amplification, and SNP details (including the flanking sequences) when relevant. QTLs link the genetic data to the phenomic data in GnpIS-coreDB and to synteny data displayed by the PlantSyntenyViewer tool [32], [33].

The accession (i.e. germplasm) and the variables (i.e. observed trait) described with dedicated ontologies are another important key data for genetic studies as they allow linking phenotype data to genetic associations or QTLs through traits and to genotype diversity data. The genetic resources stored in GnpIS-coreDB displays the unambiguous identification of the accession used (with digital object identifier - DOI) and a rich set of associated data following the MCPD (multi-crop passport descriptors, [34]) standard: a picture, synonyms, descriptors, geolocation of the sites (origin, collecting and evaluation), the collections or panels it belongs to, the holding stock centre with a link to order the accession when possible. The phenotype data includes traceability on trials with timing, like year and temporal series, location, and environment including soil and cultural practices. The phenotype and environment variables follow the crop ontologies format [35] that includes unique identifiers for each variable which are composed of a trait description (e.g. grain yield, plant height top, spike per area), a unit and a method. All these data are displayed in the GnpIS-coreDB web interface and can be downloaded in different file formats, all compliant with the MIAPPE standard [22].

### Mining and data discovery tools

To complete this already rich integrated datasets, a gene centric data warehouse, the WheatMine, has been set-up using the well-established InterMine tool [24]. The gene card displays gene function, gene ontology terms, and overlapping genomic features. WheatMine [36] provides access to IWGSC RefSeq v1.0 annotation data (genes, mRNA, polypeptides, transposable elements), polymorphisms (markers) and, through pivotal objects, to genetic data (QTL, metaQTL). It is also possible to navigate from a gene card to its position on the wheat genome browser or to relevant marker details in GnpIS-coreDB.

Figure 1 presents the concept and the tools to navigate through the key data in GnpIS.

**Figure 1.**
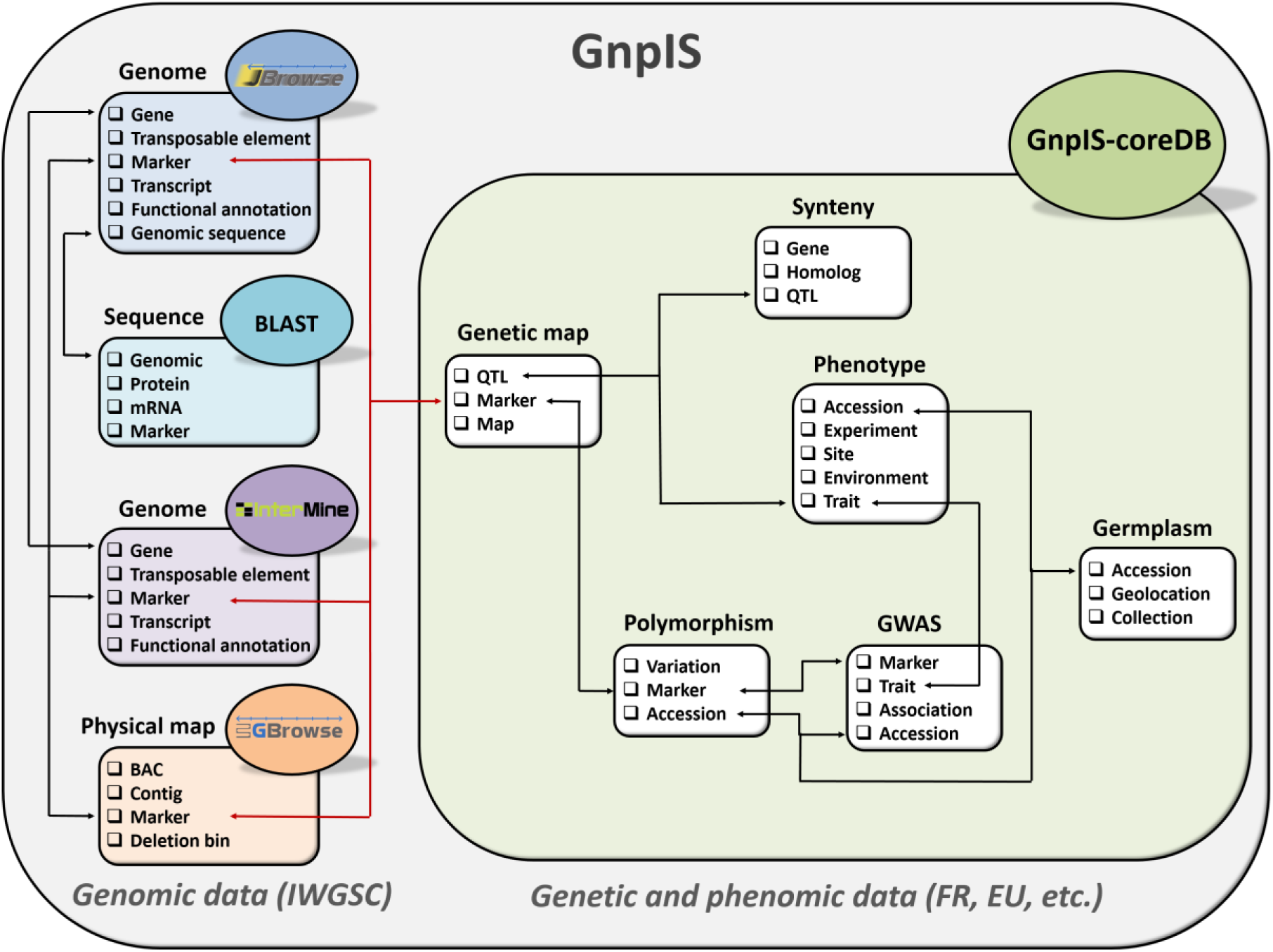
Conceptual view of wheat data links in GnpIS

Finally, to facilitate data search and access to this wealth of data, we developed a data discovery tool, which, similar to a google search, allows the user to enter keywords or terms to find all the matching information in the various data warehouses. The results are presented in a table with details on the matches (database source, type, species, description) and a direct link to the feature (e.g. a gene in a browser, a marker page in GnpIS-coreDB, etc.).

A practical use case describing how to use the Wheat@URGI portal to go from a gene sequence to find the genetic studies related is detailed in the Supplementary data.

## Conclusion and future directions

The Wheat@URGI portal hosts and gives access to essential, high quality wheat data from the IWGSC, European, and international projects. Furthermore, its added value is that it integrates different data type altogether (genomics, genetics and phenomics) and provides dedicated tools to explore them.

As new wheat resources such as GWAS, genomic selection, and pan-genome data are generated in the frame of ongoing projects, GnpIS will allow their management and integration with other data already available in the information system, linking new upcoming data to this central IWGSC genomic resource.

At a wider scale, an expert working group (EWG) of the international Wheat Initiative build an international wheat information system, called WheatIS, with the aim of providing a single-access web-based system to all available wheat data resources and bioinformatics tools [37]. The Wheat@URGI portal is a major node of the WheatIS federation that expose genomic, genetic and phenomic integrated data to the community. The WheatIS data discovery tool allows a one-stop search in GnpIS [4] (including IWGSC browsers, InterMine and GnpIS-coreDB; URGI), EnsemblPlants (EMBL-EBI) [31], CrowsNest [38] (PGSB), CR-EST [39], GBIS [40] and MetaCrop [41] (IPK), The Triticeae Toolbox (Triticeae CAP), CIMMYT DSpace and Dataverse (CIMMYT), Gramene [42] (CSH, OSU, EMBL-EBI), Cropnet (IPGPAS), WheatPan [43] (UWA) and GrainGenes [44] (USDA).

The Figure 2 presents the WheatIS ecosystem.

**Figure 2.**
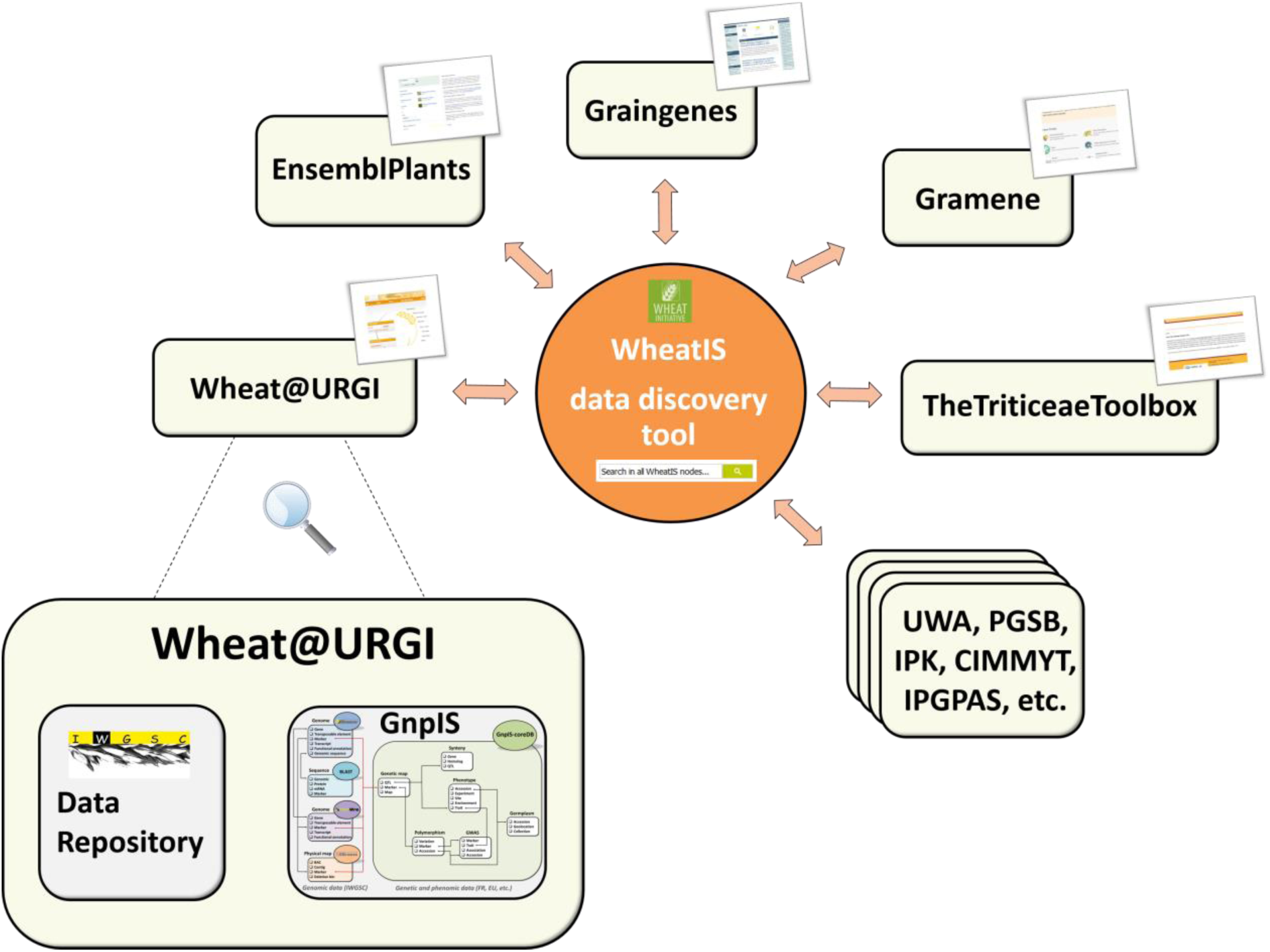
The Wheat@URGI portal node in the WheatIS ecosystem

Data integration is fundamental for researchers and breeders that want to use genomic information to improve wheat varieties. However, the diversity of data type and the concomitant lack of data harmonisation and standards hamper cross-referencing and meta-analysis. A joint action between the WheatIS EWG and a group of linked data scientists created the Wheat Data Interoperability Working Group under the Research Data Alliance (RDA) umbrella [45] to help tackle this difficult issue [46]. The Wheat@URGI portal continuously evolves its repository to follow the standard recommendations [47].

## Abbreviations

IWGSC: international wheat genome sequencing consortium
INRA: institut national de la recherche agronomique / French national institute for agricultural research
URGI: unité de recherche génomique info / research unit in genomics and bioinformatics
FAIR: findable, accessible, interoperable, reusable
BLAST: basic local alignment search tool
HICF: high-information-content fingerprinting
WGP^™^: whole genome profiling
BAC: bacterial artificial chromosome
FPC: fingerprinted contig
LTC: linear topological contig
CSS: chromosome survey sequence
POPSEQ: population sequencing
RNA: ribonucleic acid
TPM: transcripts per kilobase million
VCF: variant call format
SSR: simple sequence repeats
SNP: single nucleotide polymorphism
DArT: d iversity arrays technology
QTL: quantitative trait loci
GWAS: genome-wide association study
cM: centimorgan
HSP: high scoring pairs
PCR: polymerase chain reaction
DOI: digital object identifier
MCPD: multi-crop passport descriptors
MIAPPE: minimum information about a plant phenotyping experiment
EWG: expert working group
EMBL-EBI: European bioinformatics institute
PGSB: plant genome and systems biology group
IPK: Leibniz institute of plant genetics and crop plant research
CIMMYT: international maize and wheat improvement center
CSH: Cold Spring Harbor laboratory
OSU: Ohio State University
IPGPAS: institute of plant genetics of the Polish academy of science
UWA: University of Western Australia
USDA: U.S. department of agriculture
EWG: expert working group
RDA: research data alliance

## Declarations

### Ethics approval and consent to participate

Not applicable.

### Consent for publication

Not applicable.

### Availability of data and materials

The open access data (including all the IWGSC data) are available through the Wheat@URGI portal: https://wheat-urgi.versailles.inra.fr.

## Competiting interest

The authors declare that they have no competing interests.

## Authors’ contributions

MA, JR, TL, FA, KE designed, developed and filled the IWGSC data repository.

MA, TL, RF, FA, CP, NM, SD, EK, CM, CG, MLo, MLa, DS, AFAB, HQ designed, developed and filled the GnpIS information system.

FC, HR, PL, NG, JS, CF, IWGSC, EP generated, submitted the data and give feedback on the tools.

MA, JR, EP, KE, AFAB, HQ draft the manuscript.

All authors read and approved the final manuscript.

## Acknowledgements

The authors would like to thank for their help or advices at various stages of the project, the following people from INRA-URGI: Véronique Jamilloux, Joëlle Amselem, Dorothée Charruaud, Guillaume Cornut, Laura Burlot, Florian Philippe, Nicolas Francillonne, Loïc Couderc, Daphné Verdelet, Baptiste Brault, Kirsley Chennen; from INRA-GDEC: Jacques Le Gouis, Gilles Charmet, Fran#x00E7;ois Balfourier, Pierre Sourdille, Catherine Ravel, Fran#x00E7;ois-Xavier Oury, Audrey Didier; from INRA-DIST: Esther Dzale, Sophie Aubin, Odile Hologne; and from INRA-Agronomie: Arnaud Gauffreteau.

Thanks to Isabelle Caugant (IWGSC), Héléne Lucas (Wheat Initiative), the International Wheat Genome Sequencing Consortium and its sponsors, the WheatIS expert working group, the URGI platform, and all the data submitters.

## Supplementary data

### Software technologies

The Wheat@URGI portal website is based on eZ Publish v4 open source content management system (https://ez.no/) using the PHP language and a MySQL database (https://www.mvsql.com/).

The genome browsers are based on the GMOD (http://gmod.org/wiki/Main_Page) GBrowse v2.33 [26] and JBrowse v1.11.5 [28] built with JavaScript and HTML5. We customized GBrowse to display the physical map data. The gff3 file is generated from the .fpc file obtained by the data producer using the FPC [8] or the LTC [9] tools.

The stand-alone BLAST web interface implemented at URGI is based on ViroBLAST [48], customized to obtain a user-friendly grouping of searched databanks and visualization of the results. A robust file download system was also developed using a home-made php script to handle big data volume.

GnpIS-coreDB is a URGI development using state of the art technologies: Java EE framework (http://www.oracle.com/technetwork/iava/iavaee/overview/index.html). GWT (Google Web Toolkit, http://www.gwtproject.org/). Spring boot v1.4 (https://projects.spring.io/spring-boot/). PostgreSQL relational database v9.6 (https://www.postgresql.org/) and Elasticsearch NoSQL database v2.3.3 (https://www.elastic.co/). To set-up a GnpIS-coreDB dedicated to the wheat community, a filter allowing to display only the wheat data (*Triticum*, *Aegilops*) and barley data (*Hordeum*) was developed. This filter relies on a variable-length multidimensional arrays field in the PostgreSQL database. It is completely transparent to the user and allows him to navigate in GnpIS-coreDB through wheat data only. New versions of the GnpIS-coreDB software are deposited in the APP, the European body for protecting authors and publishers of digital works (http://www.app.asso.fr/en/welcome.html).

WheatMine uses InterMine [24] v1.8.3 which provides a fast, flexible and user friendly access to integrated data by multiple ways: a browser, a query builder and a region search tool. Users can filter their favorite features, save their own queries, and export results in many different formats (GFF3, BED or XML). An On-line documentation and pre-computed queries are also available.

The data discovery tool relies on the Solr full-text indexing technology v6.6.2 (http://lucene.apache.org/solr/). We used a restriction on the wheat and barley species to search only the corresponding data in the indexed databases. The tool was packaged and is downloadable (https://wheat-urgi.versailles.inra.fr/Proiects/Wheat-Information-System/SolR-tool-package).

### Usage Statistics

**Table S1.**
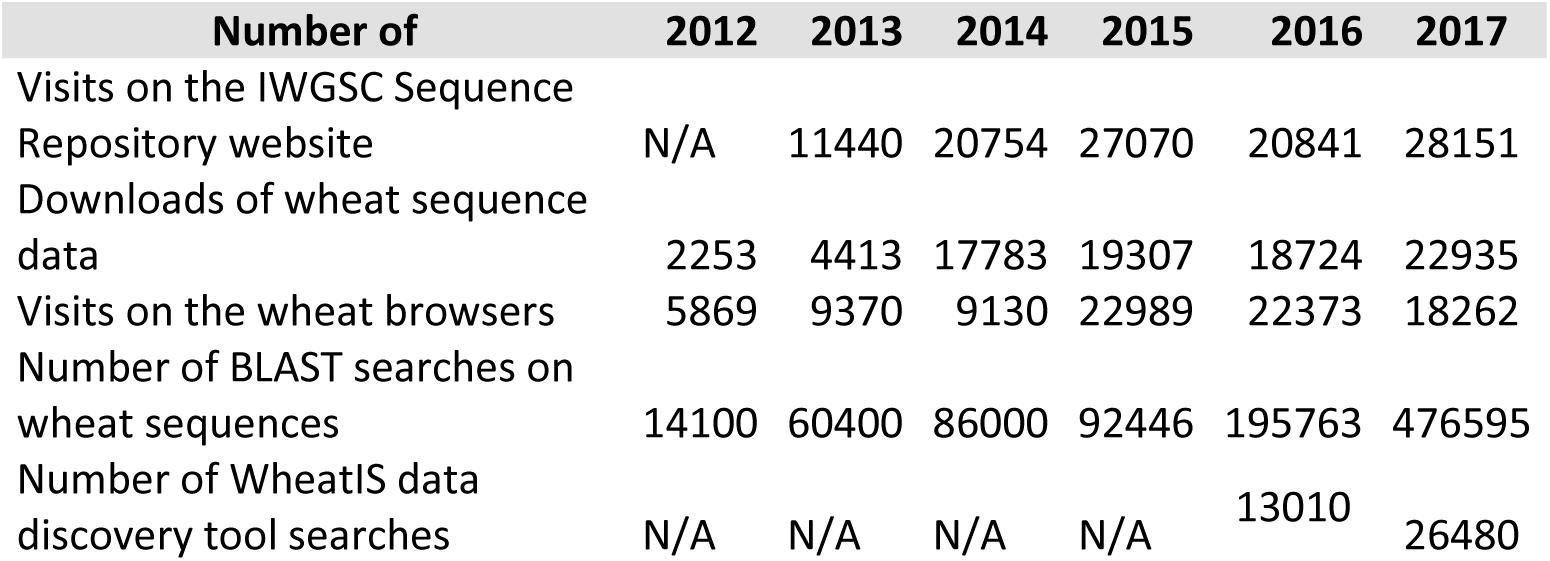
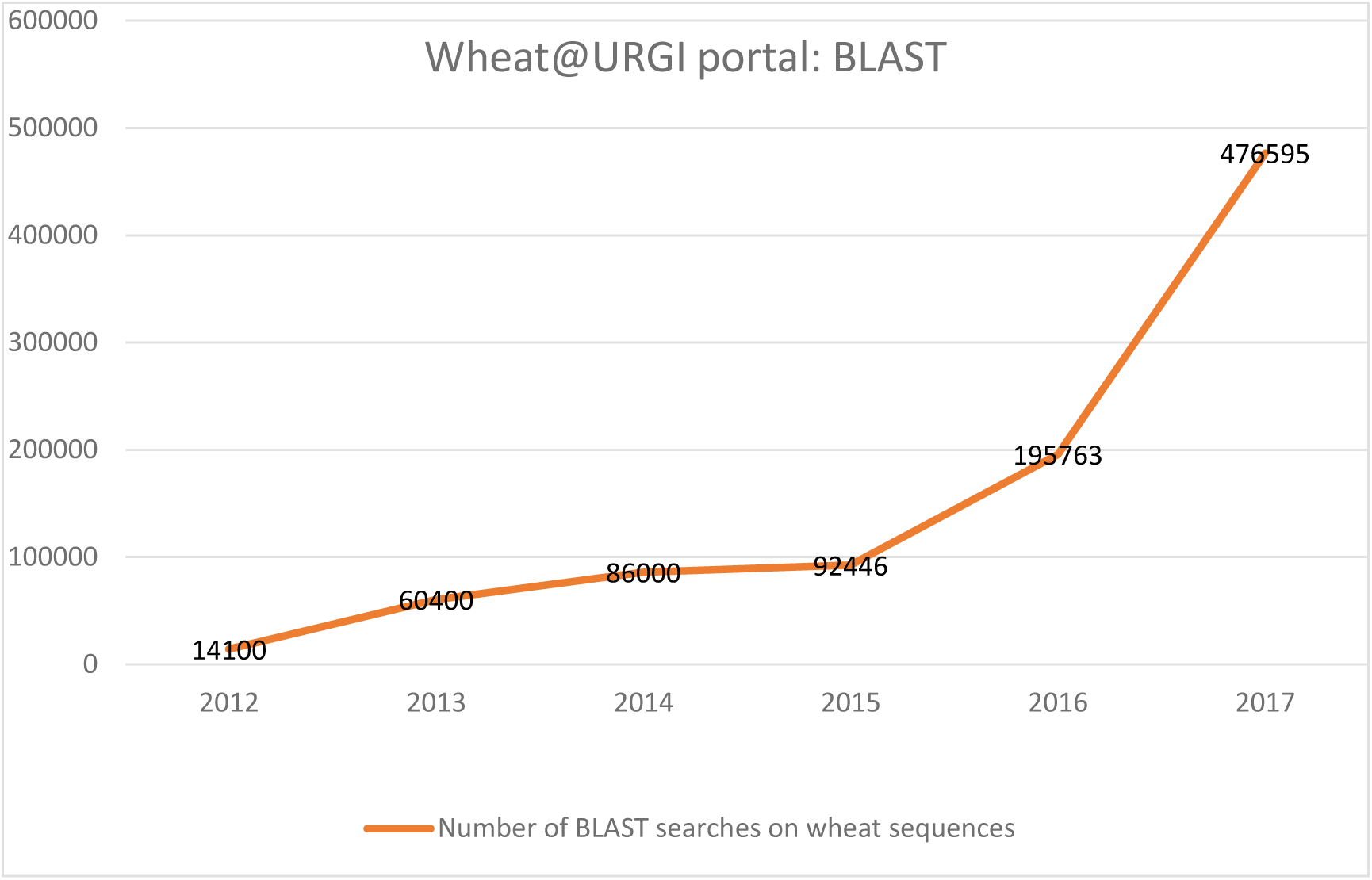
Usage statistics of genomics data in the Wheat@URGI portal (all numbers exclude web-robots and internal IP).

### Use case example

A researcher in genomics works on his wheat favorite gene. He wants to explore all the genomic data in the vicinity of this gene and find out if there are genetic studies pointing to the genomic regions where the gene is located. He searches the gene name (e.g. TraesCS5A01G033100) in the data discovery tool (https://wheat-urgi.versailles.inra.fr, Fig S1.1A) or BLAST the sequence of the gene against the IWGSC RefSeq v1.0 (https://urgi.versailles.inra.fr/blast_iwgsc/, Fig S1.1B).

**Figure S1.**
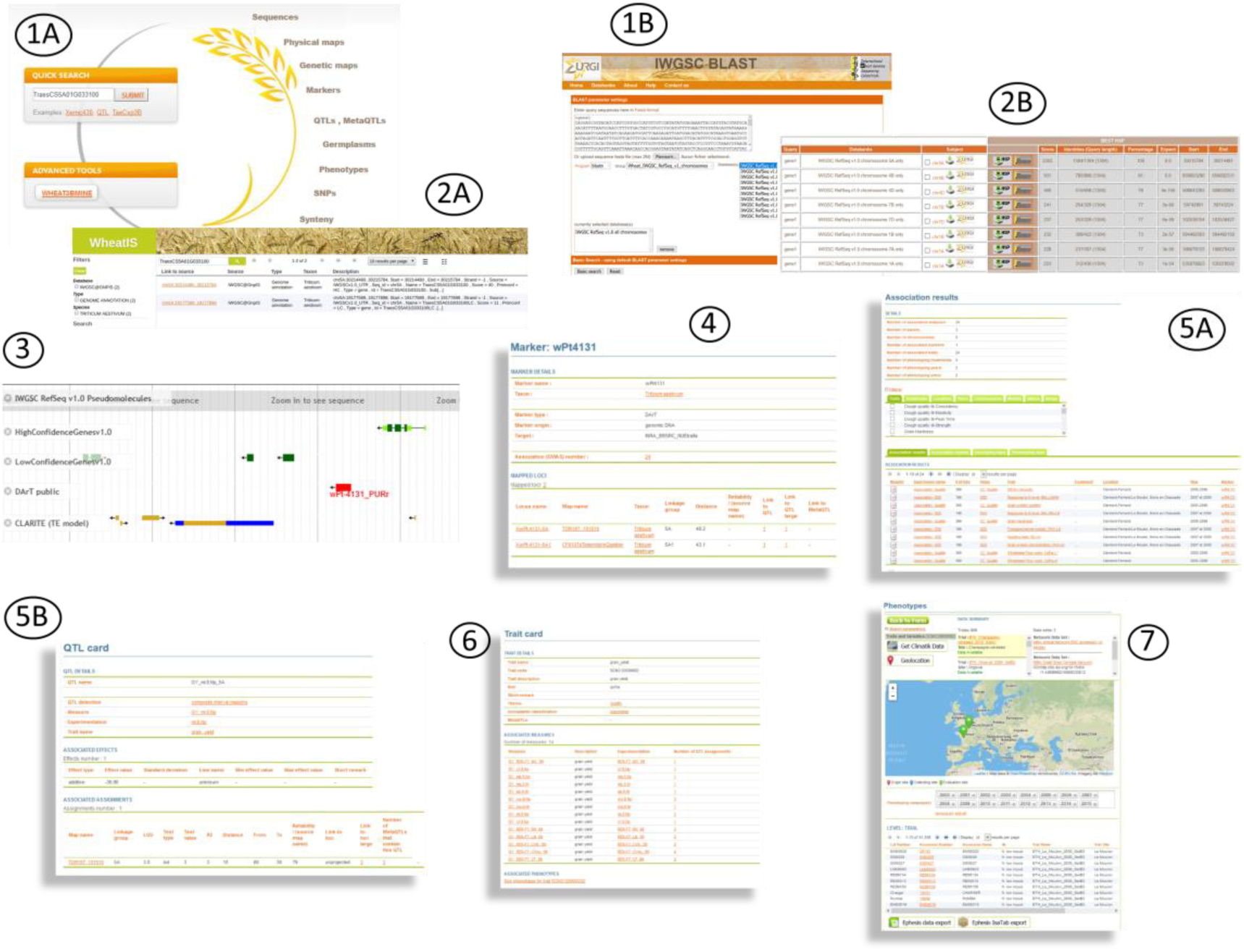
Printscreens of the web interfaces.

The results are displayed in a table (Fig S1.2A, Fig S1.2B) with links to the JBrowse directly zooming on the gene (https://urgi.versailles.inra.fr/ibrowseiwgsc/gmodjbrowse/?data=myData%2FIWGSC_RefSeq_v1.0&loc=chr5A%3A30211546..30218715&tracks=DNA%2CHighConfidenceGenesv1.0%2CLowConfidenceGenesv1.0%2CrepeatRegion%2CrepeatMasker%2CDART_PUBLIC_SUMMARY&highlight=chr5A%3A30214481..30215784%20(-%20strand)%20(TraesCS5A01G033100)). He explores the region around the gene and finds a marker (e.g. wPt-4131_PURr, Fig S1.3). By clicking on the marker, he obtains additional information stored in GnpIS-coreDB (https://urgi.versailles.inra.fr/GnpMap/mapping/id.do?action=MARKER&id=40393. Fig S1.4) showing that the marker is used in a GWAS experiments (https://urgi.versailles.inra.fr/association/association/viewer.do?results/markerIds=40393. Fig S1.5A) and is linked to a QTL (e.g. GY_ml.8.Np_5A, https://urgi.versailles.inra.fr/GnpMap/mapping/id.do?action=QTL&id=59588. Fig S1.5B). From the Trait description of this QTL (https://urgi.versailles.inra.fr/GnpMap/mapping/id.do?action=TRAIT&id=255. Fig S1.6), he displays all the phenotyping experiment performed on this trait (https://urgi.versailles.inra.fr/ephesis/ephesis/viewer.do#dataResults/traitCode=SCNO:00000002. Fig S1.7).

## References

1. IWGSC website. http://www.wheatgenome.org/. Accessed 10 April 2018.

2. IWGSC, 2018, under review.

3. Wilkinson MD, Dumontier M, Aalbersberg IJJ, Appleton G, Axton M, Baak A, et al. The FAIR Guiding Principles for scientific data management and stewardship. Sci Data. 2016;3:160018.

4. Steinbach D, Alaux M, Amselem J, Choisne N, Durand S, Flores R, et al. GnpIS: an information system to integrate genetic and genomic data from plants and fungi. Database J Biol Databases Curation. 2013;2013:bat058.

5. Wheat@URGI portal. https://wheat-urgi.versailles.inra.fr. Accessed 10 April 2018.

6. Nelson WM, Bharti AK, Butler E, Wei F, Fuks G, Kim H, et al. Whole-Genome Validation of High-Information-Content Fingerprinting. Plant Physiol. 2005;139:27–38.

7. Philippe R, Choulet F, Paux E, van Oeveren J, Tang J, Wittenberg AH, et al. Whole Genome Profiling provides a robust framework for physical mapping and sequencing in the highly complex and repetitive wheat genome. BMC Genomics. 2012;13:47.

8. Soderlund C, Humphray S, Dunham A, French L. Contigs built with fingerprints, markers, and FPC V4.7. Genome Res. 2000;10:1772–87.

9. Frenkel Z, Paux E, Mester D, Feuillet C, Korol A. LTC: a novel algorithm to improve the efficiency of contig assembly for physical mapping in complex genomes. BMC Bioinformatics. 2010;11:584.

10. French plant genomic resource centre. https://cnrgv.toulouse.inra.fr/en. Accessed 10 April 2018.

11. Altschul SF, Gish W, Miller W, Myers EW, Lipman DJ. Basic local alignment search tool. J Mol Biol. 1990;215:403–10.

12. International Wheat Genome Sequencing Consortium (IWGSC). A chromosome-based draft sequence of the hexaploid bread wheat (Triticum aestivum) genome. Science. 2014;345:1251788.

13. Choulet F, Alberti A, Theil S, Glover N, Barbe V, Daron J, et al. Structural and functional partitioning of bread wheat chromosome 3B. Science. 2014;345:1249721.

14. Mascher M, Muehlbauer GJ, Rokhsar DS, Chapman J, Schmutz J, Barry K, et al. Anchoring and ordering NGS contig assemblies by population sequencing (POPSEQ). Plant J Cell Mol Biol. 2013;76:718–27.

15. Ramirez-Gonzalez et al., 2018, submitted.

16. Jordan KW, Wang S, Lun Y, Gardiner L-J, MacLachlan R, Hucl P, et al. A haplotype map of allohexaploid wheat reveals distinct patterns of selection on homoeologous genomes. Genome Biol. 2015;16:48.

17. Rimbert H, Darrier B, Navarro J, Kitt J, Choulet F, Leveugle M, et al. High throughput SNP discovery and genotyping in hexaploid wheat. PloS One. 2018;13:e0186329.

18. Clavijo BJ, Venturini L, Schudoma C, Accinelli GG, Kaithakottil G, Wright J, et al. An improved assembly and annotation of the allohexaploid wheat genome identifies complete families of agronomic genes and provides genomic evidence for chromosomal translocations. Genome Res. 2017;

19. Luo M-C, Gu YQ, You FM, Deal KR, Ma Y, Hu Y, et al. A 4-gigabase physical map unlocks the structure and evolution of the complex genome of Aegilops tauschii, the wheat D-genome progenitor. Proc Natl Acad Sci U S A. 2013;110:7940–5.

20. GnpIS wheat data. https://wheat-urgi.versailles.inra.fr/Data. Accessed 10 April 2018.

21. Samson D, Legeai F, Karsenty E, Reboux S, Veyrieras J-B, Just J, et al. GénoPlante-Info (GPI): a collection of databases and bioinformatics resources for plant genomics. Nucleic Acids Res. 2003;31:179–82.

22. Ćwiek-Kupczyńska H, Altmann T, Arend D, Arnaud E, Chen D, Cornut G, et al. Measures for interoperability of phenotypic data: minimum information requirements and formatting. Plant Methods. 2016;12:44.

23. French small grain cereals genebank. https://www6.clermont.inra.fr/umr1095_eng/Teams/Research/Biological-Resources-Centre. Accessed 10 April 2018.

24. Kalderimis A, Lyne R, Butano D, Contrino S, Lyne M, Heimbach J, et al. InterMine: extensive web services for modern biology. Nucleic Acids Res. 2014;42:W468–472.

25. IWGSC data repository. https://wheat-urgi.versailles.inra.fr/Seq-Repository. Accessed 10 April 2018.

26. Stein LD, Mungall C, Shu S, Caudy M, Mangone M, Day A, et al. The generic genome browser: a building block for a model organism system database. Genome Res. 2002;12:1599–610.

27. GnpIS: Physical map browser. https://urgi.versailles.inra.fr/gb2/gbrowse/wheat_phys_pub. Accessed 10 April 2018.

28. Skinner ME, Uzilov AV, Stein LD, Mungall CJ, Holmes IH. JBrowse: a next-generation genome browser. Genome Res. 2009;19:1630–8.

29. GnpIS: IWGSC RefSeq v1.0 browser. https://urgi.versailles.inra.fr/jbrowseiwgsc/gmod_jbrowse/?data=myData%2FIWGSC_RefSeq_v1.0. Accessed 10 April 2018.

30. GnpIS: IWGSC BLAST tool. https://urgi.versaines.inra.fr/blast_iwgsc/blast.php. Accessed 10 April 2018.

31. Bolser DM, Staines DM, Perry E, Kersey PJ. Ensembl Plants: Integrating Tools for Visualizing, Mining, and Analyzing Plant Genomic Data. Methods Mol Biol Clifton NJ. 2017;1533:1–31.

32. GnpIS PlantSyntenyViewer. https://urgi.versailles.inra.fr/synteny/synteny/viewer.do#form/datasetId=6. Accessed 10 April 2018.

33. Pont C, Murat F, Guizard S, Flores R, Foucrier S, Bidet Y, et al. Wheat syntenome unveils new evidences of contrasted evolutionary plasticity between paleo- and neoduplicated subgenomes. Plant J Cell Mol Biol. 2013;76:1030–44.

34. Multi-Crop Passport Descriptors V.2.1. https://www.bioversityinternational.org/e-library/publications/detail/faobioversity-multi-crop-passport-descriptors-v21-mcpd-v21/. Accessed 10 April 2018.

35. Shrestha R, Matteis L, Skofic M, Portugal A, McLaren G, Hyman G, et al. Bridging the phenotypic and genetic data useful for integrated breeding through a data annotation using the Crop Ontology developed by the crop communities of practice. Front Physiol. 2012;3:326.

36. GnpIS: WheatMine tool. https://urgi.versailles.inra.fr/WheatMine. Accessed 10 April 2018.

37. WheaIS. http://www.wheatis.org/. Accessed 10 April 2018.

38. Spannagl M, Nussbaumer T, Bader KC, Martis MM, Seidel M, Kugler KG, et al. PGSB PlantsDB: updates to the database framework for comparative plant genome research. Nucleic Acids Res. 2016;44:D1141–7.

39. Künne C, Lange M, Funke T, Miehe H, Thiel T, Grosse I, et al. CR-EST: a resource for crop ESTs. Nucleic Acids Res. 2005;33:D619–21.

40. Oppermann M, Weise S, Dittmann C, Knüpffer H. GBIS: the information system of the German Genebank. Database J Biol Databases Curation [Internet]. 2015 [cited 2017 Sep 18];2015. Available from: http://www.ncbi.nlm.nih.gov/pmc/articles/PMC4423411/

41. Schreiber F, Colmsee C, Czauderna T, Grafahrend-Belau E, Hartmann A, Junker A, et al. MetaCrop 2.0: managing and exploring information about crop plant metabolism. Nucleic Acids Res. 2012;40:D1173–7.

42. Tello-Ruiz MK, Stein J, Wei S, Preece J, Olson A, Naithani S, et al. Gramene 2016: comparative plant genomics and pathway resources. Nucleic Acids Res. 2016;44:D1133–1140.

43. Montenegro JD, Golicz AA, Bayer PE, Hurgobin B, Lee H, Chan C-KK, et al. The pangenome of hexaploid bread wheat. Plant J Cell Mol Biol. 2017;90:1007–13.

44. Carollo V, Matthews DE, Lazo GR, Blake TK, Hummel DD, Lui N, et al. GrainGenes 2.0. An Improved Resource for the Small-Grains Community. Plant Physiol. 2005;139:643–51.

45. Wheat Data Interoperability Working Group of the Research Data Alliance. https://rd-alliance.org/groups/wheat-data-interoperability-wg.html. Accessed 10 April 2018.

46. Dzale Yeumo E, Alaux M, Arnaud E, Aubin S, Baumann U, Buche P, et al. Developing data interoperability using standards: A wheat community use case. F1000Research. 2017;6:1843.

47. Wheat Data Interoperability Working Group guidelines. https://ist.blogs.inra.fr/wdi/. Accessed 10 April 2018.

48. Deng W, Nickle DC, Learn GH, Maust B, Mullins JI. ViroBLAST: a stand-alone BLAST web server for flexible queries of multiple databases and user’s datasets. Bioinforma Oxf Engl. 2007;23:2334–6.

